# AuthentiCT: a model of ancient DNA damage to estimate the proportion of present-day DNA contamination

**DOI:** 10.1101/2020.03.13.991240

**Authors:** Stéphane Peyrégne, Benjamin M. Peter

## Abstract

**Summary:** Contamination from present-day DNA is a fundamental issue when studying ancient DNA from historical or archaeological material, and quantifying the amount of contamination is essential for downstream analyses. We present AuthentiCT, a command-line tool to estimate the proportion of present-day DNA contamination in ancient DNA datasets generated from single-stranded DNA libraries. The prediction is based solely on the patterns of post-mortem damage observed on ancient DNA sequences. The method has the power to quantify contamination from as few as 10,000 mapped sequences, making it particularly useful for analysing specimens that are poorly preserved or for which little data is available.

## 1 BACKGROUND

After the death of an organism, its DNA decays and is progressively lost through time [1, 2]. Under favourable conditions, DNA can preserve for hundreds of thousands of years and provide valuable information about the evolutionary history of organisms [3, 4]. Yet, only minute amounts of ancient DNA (aDNA) often remain in historical or archaeological material. In addition, most of the extracted DNA usually comes from microorganisms that spread in decaying tissues [5, 6]. Whereas microbial sequences rarely align to the reference genome used for identifying endogenous sequences if appropriate length cut-offs are used [7-9], contamination with DNA from closely related organisms represents a recurrent problem [10-12]. This is particularly true for the genomic analyses of ancient humans, as the individuals handling the specimens during excavation and at later times often leave their DNA behind [13, 14]. Because this contamination can substantially affect the results of population genetic or phylogenetic analyses, quantifying the level of contamination is crucial for downstream analyses. An estimate of the level of present-day DNA contamination is also desirable for making decisions when screening samples to identify those that can be further sequenced with reasonable effort and expenses.

Approaches to quantify the level of contamination can be divided into three categories. Some methods rely on prior knowledge of sequence differences between the contaminating and endogenous genomes [15-17]. Alternatively, if these differences are unknown *a priori*, other methods evaluate the excess of alleles compared to the expected ploidy [18-21]. A third set of methods uses patterns of chemical damage that are characteristic of aDNA [17, 22].

Among the approaches that rely on genetic differences, the most common strategy is to identify diagnostic positions expected to differ between the contaminating and endogenous sequences [16, 20]. The proportion of sequences that carry the contaminant allele at diagnostic positions represents an estimate of the level of contamination. This approach is particularly well-tailored for studying the mitochondrial genome, because it is an extensively studied non-recombining locus that is often available at high coverage. In contrast, local genealogies along the nuclear genome may differ from the overall population relationship (incomplete lineage sorting), making the identification of diagnostic positions difficult. By leveraging differences in allele frequencies between populations, it is possible to estimate the proportion of present-day human DNA contamination among nuclear sequences from archaic hominins [21-23]. However, this approach remains challenging for the analysis of early modern humans because of the lack of knowledge about rare sequence variants in the sample of interest that are unlikely to be shared with the present-day human contaminant. Thus, contamination estimates obtained from mitochondrial sequences are often used as a proxy for the level of nuclear DNA contamination [24-26]. However, the ratio of mitochondrial DNA to nuclear DNA may vary between the endogenous and contaminating DNA [27, 28], leading to potential differences in the level of contamination between the mitochondrial and nuclear genomes [29, 30].

Other approaches that compare the nuclear genetic differences between contaminating and endogenous genomes commonly exploit the ploidy of the sex chromosomes [18, 19, 31]. For instance, apparent heterozygous sites on the X chromosome of a male individual or sequences mapping to the Y chromosome for a female individual represent evidence of DNA contamination. Although these analyses do not rely on a prior knowledge about the ancestry of the ancient individual, they are either restricted to the X chromosome of male samples or cannot detect female contamination in female samples. Another concern is that the level of contamination may differ between the sex chromosomes and the autosomes if the sexes of the contaminant(s) and the ancient individual differ. Other approaches for the autosomes exist, e.g. methods using apparent alternative alleles at homozygous positions or an allelic imbalance at heterozygous positions [20, 21]. However, such approaches assume that high sequence coverage is available.

Alternatively, properties of aDNA molecules can be used to estimate contamination. Ancient DNA is typically fragmented into pieces shorter than 100bp, and exhibits miscoding base modifications that accumulate over time [32-35]. The most common miscoding lesions observed in aDNA are the results of cytosine deamination [36-39] that converts cytosine (C) into uracil (U), which is then misread as thymine (T), or 5-methylcytosine into thymine. These apparent C-to-T substitutions occur preferentially toward the ends of sequences [39], likely because single-stranded overhangs, which are common in aDNA, exhibit a rate of cytosine deamination about two orders of magnitude higher than double-stranded regions [1, 40]. To estimate present-day DNA contamination, these properties need to be formalized in a model of aDNA damage. The simplest approach (“conditional substitution analysis”) is based on a model that assumes independence between C-to-T substitutions at both ends of sequences. Testing whether these substitutions are correlated between ends may reveal a set of undamaged sequences, which are likely contaminants [3]. This method works even for low sequence coverage, but is primarily used to indicate the presence of contamination. Other methods extend this approach by considering the distribution of C-to-T substitutions along sequences, either assuming a parametric ([39], PMDtools [41] and mapDamage [42, 43]) or empirical distribution of these substitutions along sequences (contDeam [17] and aRchaic [44]). Notably, these methods assume that C-to-T substitutions in the aDNA sequences are independent from each other.

Here, we introduce a novel model for aDNA damage that does not assume independence between C-to-T substitutions. Our implementation, AuthentiCT, allows both estimation of the present-day DNA contamination rate and deconvolution of endogenous and contaminating sequences solely based on patterns of aDNA damage. Applying this method to both simulated and existing aDNA datasets, we find that present-day DNA contamination can be estimated from as few as 10,000 sequences, making it a practical tool in the screening of samples for aDNA preservation.

## 2 RESULTS

### 2.1 Method overview

In this section, we first motivate our approach by studying the features of aDNA damage. We then formalize a model of aDNA damage, and develop a mixture model to describe and distinguish endogenous from putative contaminating sequences.

#### 2.1.1 Ancient DNA deamination patterns used in this study

Deamination patterns in aDNA sequences depend on the DNA library preparation method [45]. Some methods involve ligation of adapters to double-stranded DNA (“double-stranded libraries”, [46]) while other methods convert the two DNA strands into separate library molecules (“single-stranded libraries”, [47]). Here, we focus on the damage patterns in single-stranded libraries, as they fully preserve the strand orientation of the sequenced DNA fragments and are widely used in aDNA studies [48-54].

While C-to-T substitutions occur predominantly at the ends of DNA fragments, they are also common in the internal part [39, 58]. Our model is motivated by the finding that these internal C-to-Ts are not independent from each other (**Figure 1**). Excluding the first and last five bases to mask potential overhangs, we found that C-to-Ts are particularly common in adjacent positions in many samples, with a significant deviation from the geometric distribution expected from independent events (p<10^−15^, chi-square goodness-of-fit test; see Supplementary Note 1 for more details or for results excluding the first and last ten bases), and from a control using sheared present-day human DNA that was treated like aDNA [9]. Single-stranded regions inside aDNA fragments represent a possible cause.

**Figure 1:**
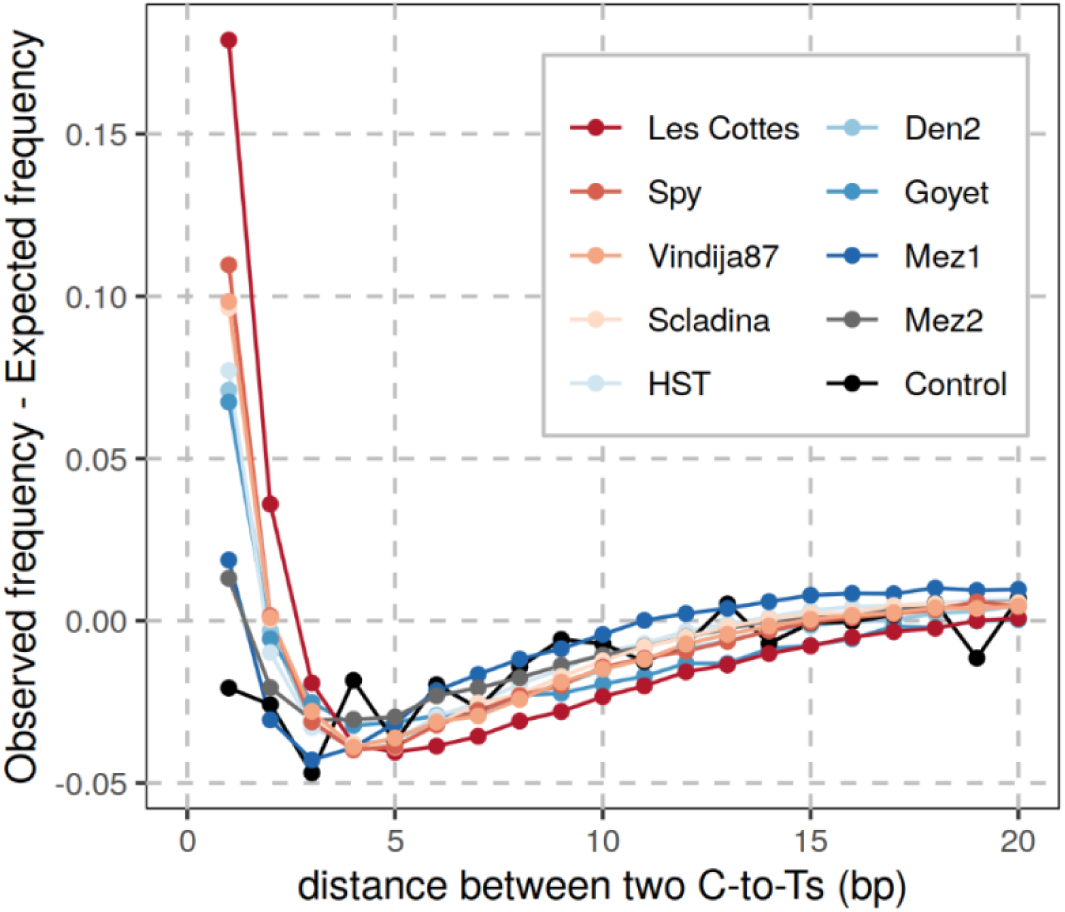
Excess of directly adjacent C-to-T substitutions inside aDNA sequences. The observed and expected frequencies of the distance between pairs of C-to-T substitutions are compared among sequences that exhibit two internal C-to-T substitutions (excluding the first and last 5 bases). The colours correspond to sequence data from different archaic humans (>500,000 sequences each; Les Cottés Z4-1514, Spy 94a, Vindija 87, Mezmaiskaya 2, Goyet Q56-1 from [55]; Scladina I-4A and Hohlenstein-Stadel from [23]; Denisova 2 from [56]; Mezmaiskaya 1 from [57]). The present-day human DNA control dataset is represented in black [9]. All sequences derived from single-stranded DNA libraries.

#### 2.1.2 Model of ancient DNA damage

Motivated by this finding, we developed a model of aDNA damage that jointly models all C-to-T substitutions, accounting for the observed clustering of C-to-T substitutions within a sequence. We used a Hidden Markov Model (HMM), where each potentially deaminated site in the reference is an informative site, i.e. Cs or Gs for sequences aligning to the forward or reverse strand, respectively. Other positions will give the same likelihood for the endogenous and contaminating DNA models, and are therefore ignored in both models. At the C or G positions of the reference genome, we classify observations either as emitting “M” (matches the reference allele), “D” (differs from the reference allele; compatible with a deamination) or “E” (other mismatches corresponding to sequencing errors or mutations) (**Figure 2**).

**Figure 2:**
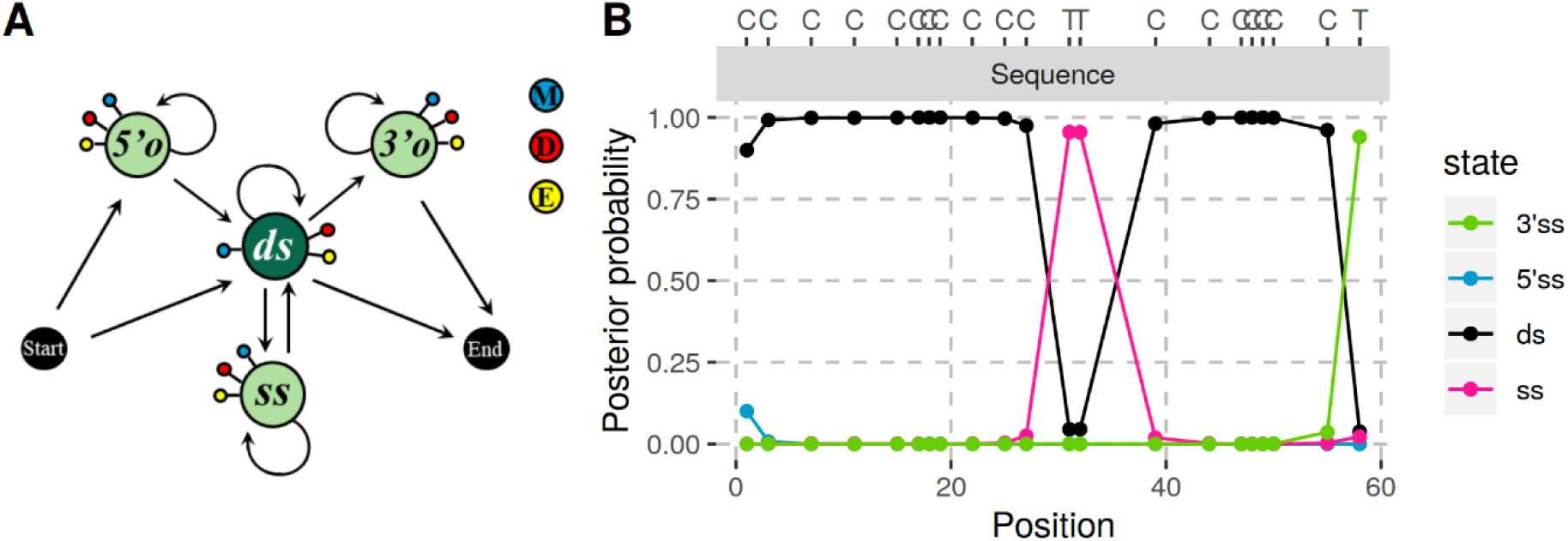
Graphical representation of the model (A) and illustration of the posterior decoding (B). (A) States are depicted by nodes and transitions by edges. Each state emits a match to the reference M (blue), or a mismatch, which can either be compatible with cytosine deamination, D (red), or an error (or polymorphism), E (yellow). Single-stranded states (*5’o, 3’o* and *ss*) and the double-stranded state (*ds*) are in light and dark green, respectively. (B) The posterior probability for each state is shown with different colors. We show bases on top for positions in the sequence (grey bar) that align to a C in the reference. See Supplementary Note 2 for an evaluation of the posterior decoding.

The model has four hidden states corresponding to double-stranded or single-stranded stretches, and 5’ and 3’ single-stranded overhangs, which we denote as *ds, ss, 5’o* and *3’o* states, respectively. At the first position of the sequence alignment, the chain starts in a 5’ single-stranded overhang with the probability *o*, or in a double-stranded state with the probability 1-*o*. Then, the lengths of the different regions follow geometric distributions, with the parameters *l*_*o*_, *l*_*ss*_ or *l*_*ds*_ for the overhangs, single-stranded and double-stranded regions, respectively. We therefore assumed the following matrix of transition probabilities:

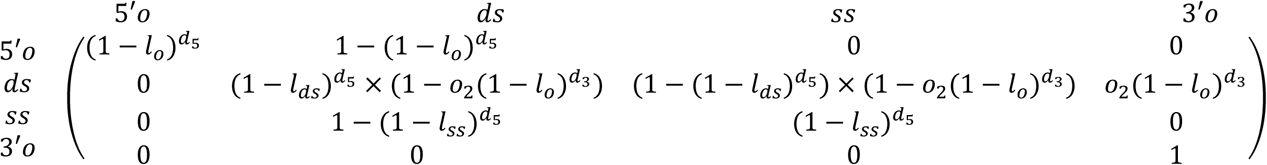

with *d*_*5*_ representing the distance from the previous observation (set to the current position for the initial observation), *d*_*3*_ the distance to the end of the sequence, and *o*_*2*_ the probability of a 3’ single-stranded overhang. The modelling of the state transitions between informative sites assumes that only one transition happens between two informative sites.

The chain ends with the following transition probabilities:

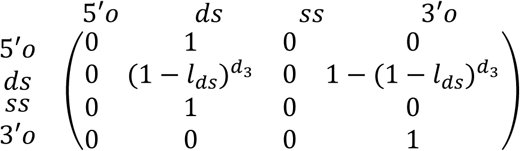

For the emission probabilities of the three possible observations mentioned above, we assumed that all single-stranded states (*5’o, 3’o* and *ss*) have emissions:

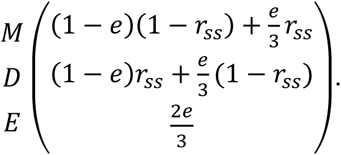

Similarly, the emission vector for the double-stranded state is:

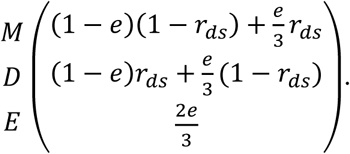

Here, *r*_*ss*_ and *r*_*ds*_ denote the deamination rates in single-stranded regions (including the single-stranded overhangs) and double-stranded regions, respectively. We model polymorphisms and errors using a single rate, *e*, as these are indistinguishable without prior knowledge of polymorphic sites in the genomes of the source populations/species. We also note that all types of substitutions are assumed to be equally likely, a simplification that has also been used in previous work [15, 18].

#### 2.1.3 Model of present-day DNA contamination

To identify contamination, we contrast the aDNA model with a model for DNA without deamination. We assumed that any difference to the reference genome arose from a constant mismatch rate *e* along the sequence. Assuming independence between sites, the probability of the data is simply: *e*^*d*^(1 − *e*)^*s*^ where *d* and *s* are the number of mismatches and matches to the reference at informative positions, respectively. We used the same rate *e* for both endogenous and contaminating sequences.

#### 2.1.4 Estimating present-day DNA contamination

We use a mixture model to estimate the overall contamination rate *c*. We denote the *i*^*th*^ sequence as X_i_, assuming we have *N* sequences in total. Using the aDNA sequence model, for each sequence, we calculate P_E_(X_i_ | *θ, e*), the probability of the sequence given that the corresponding DNA fragment is endogenous, conditional on the HMM parameters *θ*. Similarly, using the model of contaminating DNA outlined above, we calculate P_C_(X_i_ | *e*), the probability of the sequence given that it originates from a contaminating DNA fragment. Therefore, we have P(X_i_ | *θ, e*) = *c* P_C_(X_i_|*e*) + (1-*c*) P_E_(X_i_ | *θ, e*). Further assuming sequences are independent, the complete likelihood is 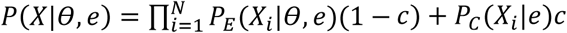. We obtain a maximum likelihood estimate of *c* using the L-BFGS-B algorithm (tolerance: 10^−10^) implemented in scipy.optimize (version 1.3.1), and estimate standard errors using the Hessian matrix of the likelihood function, generated using the numdifftools library (version 0.9.39). Assuming that the maximum likelihood estimates are normally distributed, the 95% confidence intervals (CI) are approximated as ± 1.96 standard errors.

### 2.2 Evaluation of AuthentiCT

We implemented this model in a program called AuthentiCT. In this section, we evaluate how well AuthentiCT is able to estimate the proportion of contaminating sequences, and to separate aDNA from present-day DNA sequences.

#### 2.2.1 Inference of present-day DNA contamination rates

##### Assessing the accuracy with artificial mixtures of ancient and present-day DNA

To test if AuthentiCT can estimate the proportion of present-day DNA contamination, we created artificial mixtures of aDNA and present-day DNA sequences in varying proportions, from 5% to 95%, in steps of 5%. As present-day contaminant, we used sequences generated from present-day human DNA previously sheared to short fragments and treated like aDNA (mimicking the treatment of a genuine present-day DNA contamination, [9]). As aDNA sequences, we used sequences from archaic datasets generated from single-stranded libraries that exhibit minimal amounts of present-day human DNA contamination [55, 57].

For each dataset, we then compared the contamination rate estimates from AuthentiCT to the estimates from contDeam [17] and from the conditional substitution analysis ([22], as computed in [23]) (**Figure 3**). While the latter approach underestimates the contamination proportion (average bias: - 6.73%; Root Mean Square Error (RMSE): 0.0741; based on 100,000 sequences), contDeam overestimates it (average bias: 2.36-1.19%; RMSE: 0.0396-0.0320; based on 10,000 and 100,000 sequences, respectively). In contrast, our method yields more accurate estimates (average bias: 0.05-0.42%; RMSE: 0.0194-0.0191; based on 10,000 and 100,000 sequence, respectively). We note some variability in the results depending on the dataset used, which may reflect different properties that are not under our control (e.g. additional contamination in the Neandertal datasets or differences in error rates between the present-day and ancient DNA sequences).

**Figure 3:**
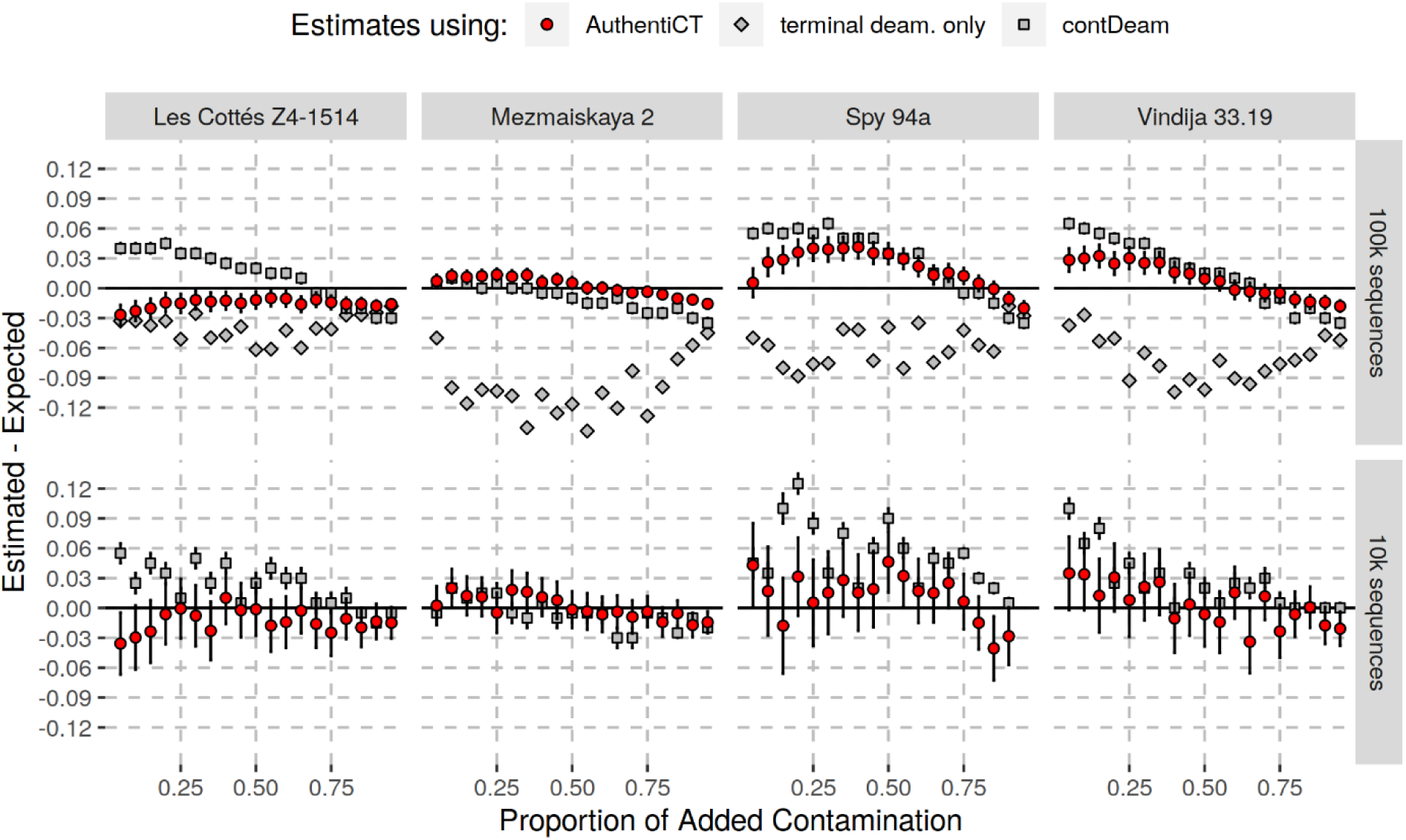
Contamination estimates on artificial mixtures of Neandertal and present-day human DNA sequences. Estimates from AuthentiCT are shown with red dots, whereas the estimates from two other methods are shown in grey [17, 22]. Each point represents the difference between the estimated proportion of contamination and the proportion of present-day human DNA sequences introduced in the corresponding Neandertal dataset (different panels). Error bars correspond to 95% CI. There were not enough observations with 10,000 sequences for the method relying only on terminal deamination [3]. We note that the Neandertal datasets, particularly Spy 94a, may contain some present-day human DNA contamination in addition to the human contamination we introduced (see Supplementary Table S1 for the libraries used and the associated contamination estimates).

##### Exploring the limits of the method with simulations

To further evaluate AuthentiCT in scenarios where we have full control over the parameters, we simulated aDNA and present-day DNA sequences using the model described above, varying deamination rates, error rates, sequence lengths, GC contents, and numbers of sequences. Unless stated otherwise, each dataset contained 100,000 sequences with lengths following a shifted geometric distribution with minimum and mean lengths of 35 and 45 bp, respectively. By default, we use a GC content of 40%, a terminal deamination rate of 0.5 and an error/divergence rate of 0.001, and set the HMM-parameters to *o*=0.5, *o*_*2*_=0.5, *l*_*o*_=0.34, *l*_*ss*_=0.20 and *l*_*ds*_=0.003.

We found that AuthentiCT performs well for datasets of 10,000 or more sequences (RMSE: 0.010 and 0.009; average bias: 0.006 and 0.007; **Figure 4A**) and its performance is consistent over a wide range of deamination rates (from 0.03 to 0.5 in **Figure 4B**; RMSE between 0.005 and 0.013), albeit with larger confidence intervals for lower deamination rates. The least reliable estimates were obtained for small datasets (1,000 sequences) with low contamination rates (RMSE: 0.028; average bias: 0.010; **Figure 4A**). We note that AuthentiCT overestimates contamination for low contamination rates (average bias of 0.020 and 0.009 for contamination estimates below 0.25 with a terminal deamination rate of 0.15 and 0.5, respectively). This likely represents overfitting to short sequences with few informative sites, as the bias decreases with longer sequences or higher GC contents (Supplementary Figure S1).

**Figure 4:**
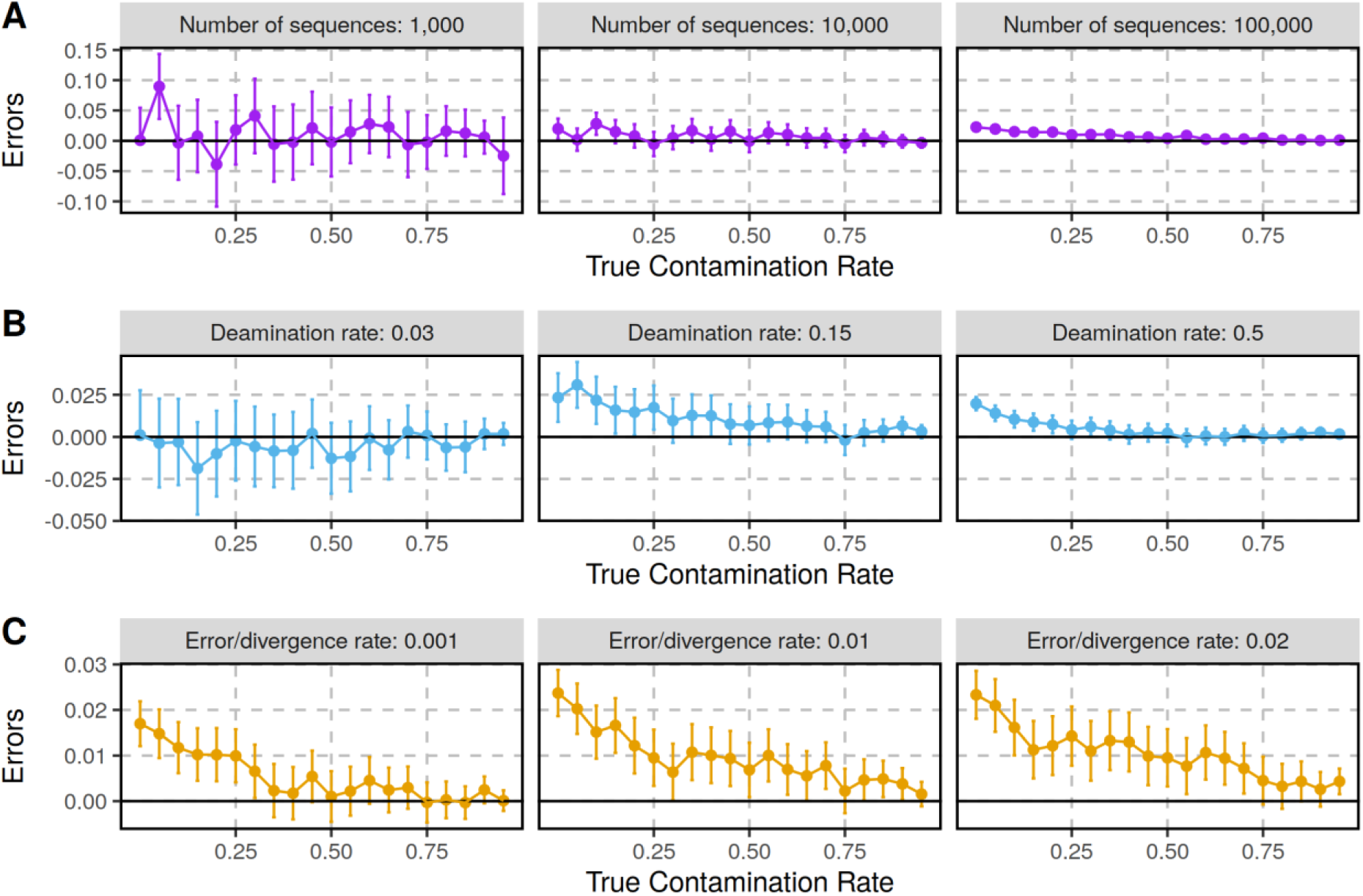
Contamination estimates on simulated datasets. The datasets differ in the number of sequences **(A)**, deamination rates **(B)** and sequencing error/divergence rates **(C).** Each point corresponds to a set of simulated sequences with varying proportions of contamination (x-axis) from 0% to 95% in steps of 5%. The errors (y-axis) correspond to the difference between the estimated and true contamination rates. The bars correspond to the 95% CI.

Another variable that may affect contamination estimates is the rate of C-to-T substitutions to the reference genome that are not induced by deamination. Although the sequencing error rate would typically not exceed 1% on sequencing platforms commonly used for ancient DNA sequencing [59, 60], divergence to the reference genome may be an issue. We therefore tested our method on simulated sequences with varying substitution rates, assuming the same rate for endogenous and contaminating sequences (see Supplementary Figure S2 for results with different rates). As expected, an increase of the substitution rate leads to a decrease in accuracy of the contamination estimates (RMSE of 0.006, 0.010 and 0.011 for substitution rates of 0.001, 0.01 and 0.02, respectively; average bias of 0.005, 0.009 and 0.01 for the same substitution rates; **Figure 4C**). We note that divergence can also lead to alignment issues that may further introduce biases in the damage patterns, as mismatches to the reference may lead to the selective loss of sequences with additional C-to-T substitutions.

##### Application to real ancient DNA datasets

To further validate the method, we next applied AuthentiCT to published sequences from archaic human specimens [22, 23, 55, 57] without introducing sequences from present-day DNA. We compared our results with a previous approach (Peyregne et al 2019) that relies solely on the divergence to a present-day African genome (HGDP00456, [45]). This independent method works well for Neandertals (as the contaminating modern human DNA will be much more similar to the African genome than the Neandertal genome), but will not translate to early modern humans. Here, we use the F(A|B) statistic as a measure of divergence, as it varies little between individuals from the same population [61]. The value of this statistic for a contaminated sample is simply a linear combination of the values for the endogenous and contaminating genomes, i.e. F_observed_ = *c* × F_contaminant_ + (1 − *c*) × F_endogenous_ where *c* is the contamination rate. We set F_contaminant_ to 0.275 (computed from the genotype calls of a present-day European genome, HGDP00521 [45]) and F_endogenous_ to 0.176 (Table S20 in [57]).

We note, however, that the two approaches measure slightly different contamination proportions. While AuthentiCT relies on every sequence, F(A|B) relies only on sequences that overlap informative positions. Therefore, AuthentiCT yields contamination estimates corresponding to the proportion of contaminating sequences, whereas the approach based on F(A|B) provides estimates corresponding to the proportion of bases that come from contaminating sequences. These estimates can differ if the length distributions of endogenous and contaminating sequences differ, as is the case for some datasets (Supplementary Note 3). To get comparable estimates of contamination, we ran AuthentiCT on the sub-set of sequences that overlap the informative sites used for quantifying contamination based on the F(A|B) statistic (**Figure 5**; see Supplementary Table S2 for the contamination estimates per sequences). Using the same sequences may also account for potential differences in the proportion of contamination along the genome. The contamination estimates from both methods are highly correlated (Pearson’s coefficient: 0.98; p-value<10^−15^; **Figure 5**).

**Figure 5:**
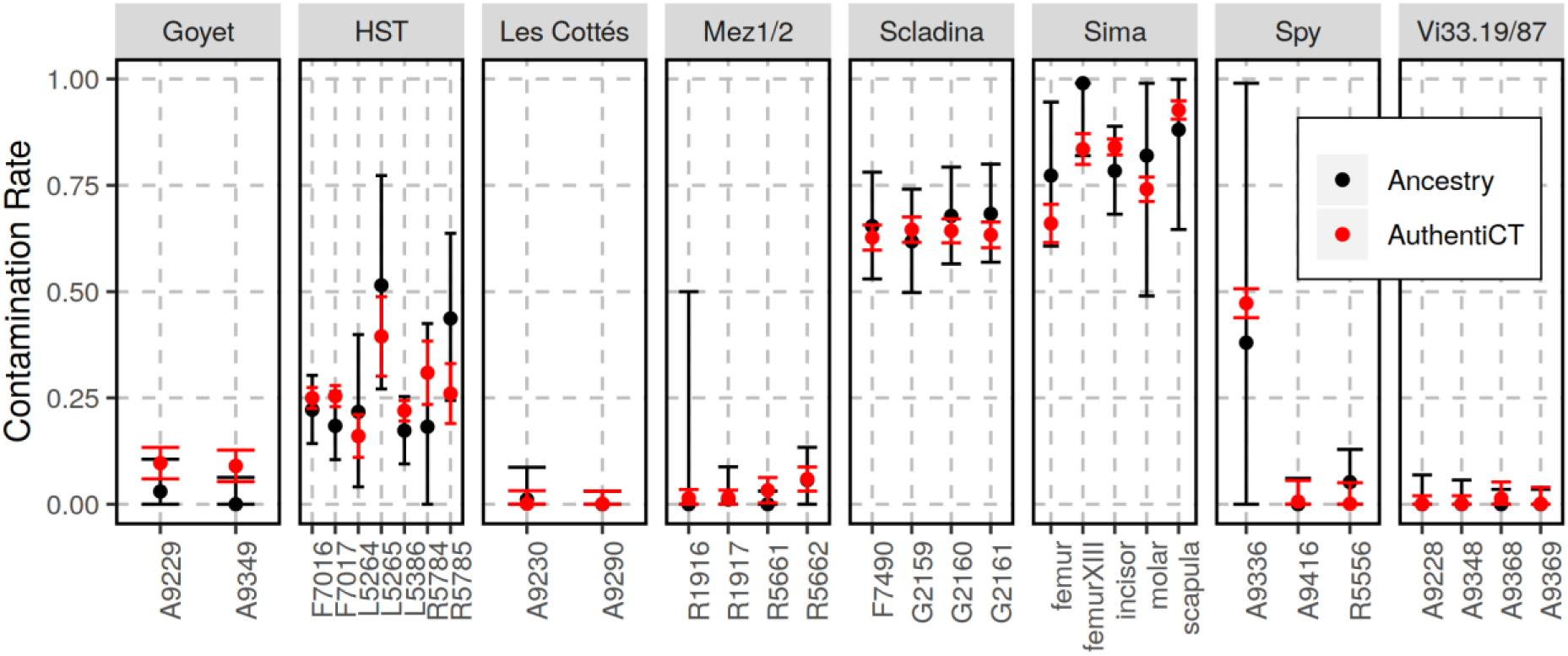
Contamination estimates in archaic DNA datasets. Estimates from AuthentiCT (in red) and a method based on the proportion of shared derived alleles with a present-day human genome (in black; see Material and Methods). When available, we used 10,000 sequences from public datasets generated from single-stranded libraries [22, 23, 55, 57]. For more details about the libraries (identifiers on the x-axis) see Supplementary Table S2. Bars represent 95% CI.

#### 2.2.2 Separating ancient DNA and present-day DNA sequences

As most downstream applications assume the absence of contamination, it is often necessary to identify endogenous sequences in contaminated aDNA samples. A common way to achieve this is to compute a likelihood-ratio for each sequence whether it is endogenous or a contaminant, as done by PMDtools [41]. Here, we investigate how the likelihoods of AuthentiCT contrasts with the ones from PMDtools (using recommended parameters) on the same artificial mixtures of Neandertal and present-day human DNA sequences described above.

A strict filter for sequences with at least one C-to-T substitution within the first or last three positions leads to about 30% of endogenous sequences being detected, with 0.4% of false positives (**Figure 6A**). At the same false positive rate, both PMDtools and AuthentiCT detect around 50% of sequences, and the likelihood models allow fine-tuning of precision with recall (see Supplementary Figure S3 for the performance on other datasets). The performance of AuthentiCT and PMDtools is similar at low false positive rates (< 0.02), which are most important for classifying sequences. However, AuthentiCT performs better for more ambiguous sequences and yields higher likelihoods for a contaminant origin for present-day DNA sequences (Supplementary Figure S4), which explains why it performs well for estimating contamination (**Figure 3**). The two methods mostly differ in their classification of sequences that exhibit only one C-to-T substitution (**Figure 6B**) in the internal part of sequences (**Figure 6C**). Compared to AuthentiCT’s classification, there is an excess of sequences classified as ancient by PMDtools that exhibit non-deaminated bases adjacent to the C-to-T substitution (**Figure 6D**). Some of these are therefore likely to represent polymorphisms or sequencing errors rather than deamination, indicating that AuthentiCT is more conservative when classifying these sequences as endogenous.

**Figure 6:**
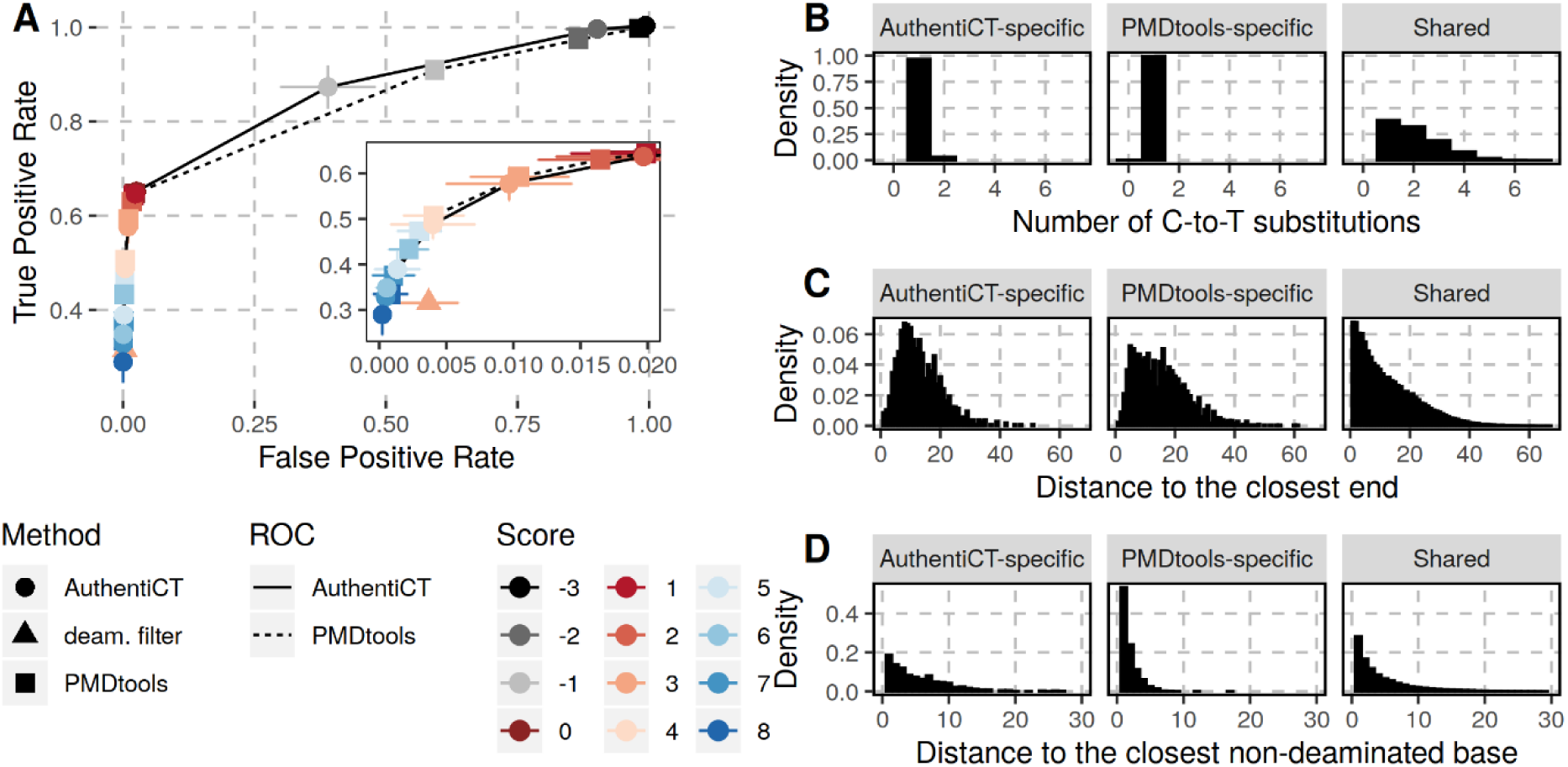
Classification of aDNA and present-day DNA sequences. **(A)** The Receiver Operating Characteristic (ROC) curves illustrate the performance of AuthentiCT (solid) and PMDtools (dashed) to identify aDNA sequences. A sequence is considered ancient if the log-likelihood ratio (score) of an ancient versus present-day origin is equal to or higher than a threshold (different colours). Each point represents the average performance over 19 datasets (of 10,000 sequences each) with varying proportions of ancient and present-day DNA sequences (5% to 95% in steps of 5%) for AuthentiCT (circles), PMDtools (squares) and a filter for sequences exhibiting at least one C-to-T substitution within the first or last three positions (deam. filter; triangles). Sequences are from Mezmaiskaya 2 (libraries A9180, A9288, A9289 and R1917 [55]) and the present-day human control. The bars correspond to two standard errors. **(B, C and D)** The distributions illustrate the number of C-to-T substitutions per sequences **(B)**, the distance between a C-to-T substitution and the closest end of the sequence **(C)** or the closest non-deaminated base **(D)** in sequences classified as ancient only by AuthentiCT (left), only by PMDtools (middle), or both (right), using a score threshold of 3 as recommended for PMDtools [41].

## 3 DISCUSSION

Estimating present-day DNA contamination in aDNA datasets remains a difficult task, particularly if the contaminating DNA is closely related to the DNA of the studied organism. Most approaches rely on genetic differences between the endogenous and contaminating genomes, which are often unknown *a priori*. Besides, the dependence on the same information used in downstream analyses is not desirable as contamination may confound signals of interest (e.g. modern human ancestry in a Neandertal genome may look like present-day human DNA contamination). Here, we developed an alternative method to estimate the proportion of present-day DNA contamination based solely on a model of aDNA damage.

AuthentiCT overcomes several shortcomings of previous methods for estimating contamination that are based on aDNA damage. Most notably, it uses every position that is potentially informative, irrespective of whether it is close to the end of a sequence or not, and accounts for clusters of C-to-T substitutions in the internal part of aDNA sequences. The latter represent a feature of aDNA deamination patterns that, to our knowledge, has not been described or used before. We demonstrated that this more detailed modelling of the distribution of C-to-T substitutions along aDNA sequences leads to more accurate estimates of the proportion of present-day DNA contamination than previous approaches. However, the performance of AuthentiCT and PMDtools to classify sequences with multiple C-to-T substitutions were almost identical, and filtering for aDNA sequences will yield very similar results for both methods. Thus, the improvement stems largely from a more confident detection of contaminant sequences. In contrast to PMDtools, a sequence with many non-deaminated bases is considered to be more likely a contaminant under the AuthentiCT model (Supplementary Figure S4).

It is important to note potential caveats. First, we assume the absence of significant levels of deamination in the contaminating DNA. This assumption does not always hold true (e.g. [13, 23, 56]) and would lead to an underestimate of the proportion of contamination (Supplementary Figure S5). One could test this by first identifying sequences that carry contaminant alleles at diagnostic positions in the mitochondrial genome and then checking for the presence of C-to-T substitutions [62]. Second, there may be populations of DNA fragments with different rates of damaged bases, even within a single sample, because of microstructural differences in DNA preservation, or because of different treatments [63]. These differences may lead to an overestimate of the proportion of present-day DNA contamination, as it would lead to an excess of sequences without C-to-T substitutions. Yet, among the 31 libraries that we tested (from 14 specimens), the contamination estimate of AuthentiCT fell only once outside the confidence intervals of another estimate from an independent method (library A9349 from the Goyet Q56-1 Neandertal, **Figure 5**). As AuthentiCT can run on as few as 10,000 sequences (in 3-10 minutes, see Supplementary Figure S6), one could split the data by sequencing run, sequence length or other covariates to obtain stratified contamination estimates. Finally, AuthentiCT is not applicable to libraries generated after treatments that alter deamination patterns, e.g. uracil-selection or treatment with a uracil-DNA-glycosylase (UDG) [64, 65].

In the future, it will be useful to extend its application to data where patterns of DNA damage differ, such as in double-stranded aDNA libraries [46]. In addition, contamination estimates might be further improved if additional features typical of aDNA are incorporated into the model, such as fragment length or the frequency of purines in the reference genome at positions flanking the sequence alignments [39].

## CONCLUSIONS

AuthentiCT is useful for estimating contamination in small datasets, e.g. when screening ancient specimens with shallow sequencing depth or when the samples are badly preserved. The independence of the contamination estimates from genetic differences between the contaminating and endogenous genomes makes this method both particularly valuable for the study of ancient human samples and broadly applicable to other species.

## DECLARATIONS

### Ethics approval and consent to participate

Not applicable.

### Consent for publication

Not applicable.

### Availability of data and materials

An open-source implementation of AuthentiCT in python and a test dataset are available on GitHub https://github.com/StephanePeyregne/AuthentiCT, under the GPLv3.0. The analysed data were generated in previous studies or are available upon reasonable request.

### Authors’ contributions

SP conceived the study. SP and BMP designed the experiments. SP implemented the software. SP performed the analyses. SP and BMP wrote the manuscript. Both authors read and approved the final manuscript.

## Acknowledgments

We thank Johann Visagie for valuable input in the implementation of AuthentiCT, Alba Bossom Mesa, Cesare de Filippo, Mateja Hajdinjak, Janet Kelso and Matthias Meyer for constructive comments on the manuscript, Svante Pääbo, Kay Prüfer and Viviane Slon for helpful discussions.

