## Supplementary Material for "AuthentiCT: a model of ancient DNA damage to estimate the proportion of present-day DNA contamination"

### Figures

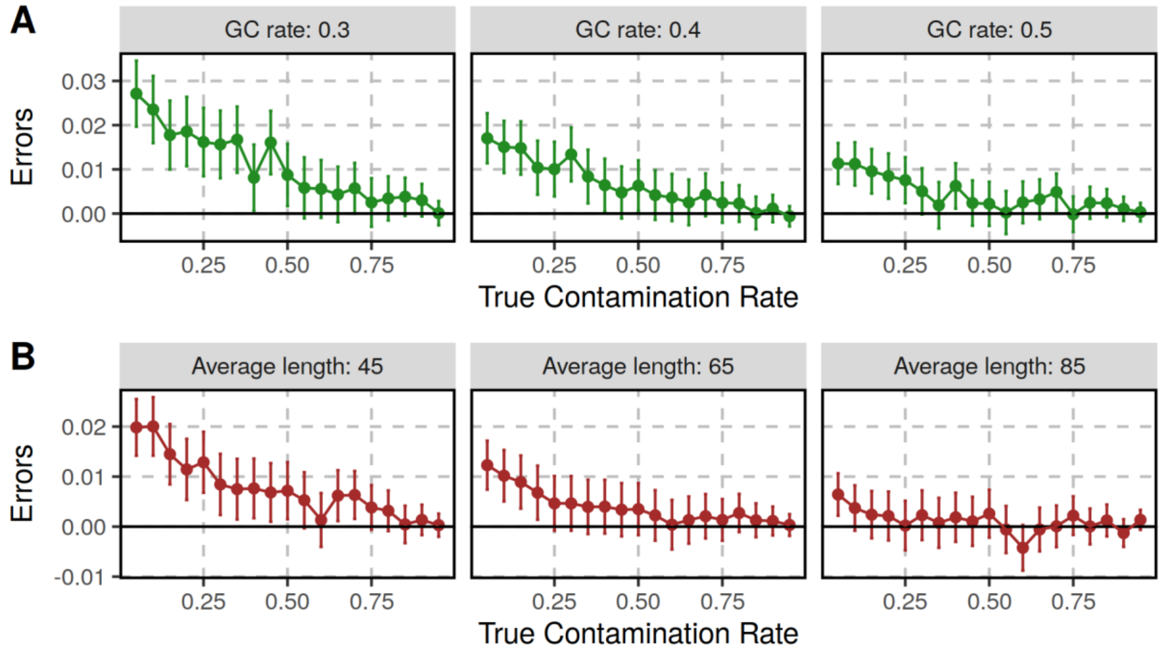

**Figure S1 : Contamination estimates on simulated datasets with different GC contents (A) and average sequence lengths (B).** Each point corresponds to the estimate for a set of 100,000 simulated sequences with varying proportions of contamination (x-axis) from 5% to 95% in steps of 5%. We set the minimum length of simulated sequences to 35 bp, the terminal deamination rate to 0.5 and 0 for the ancient and present-day DNA sequences, respectively, and the error/divergence rate to 0.001. The errors (y-axis) correspond to the difference between the estimated and true contamination rates. We note that the mean GC content of the human genome is 41% [1] and that the length of ancient DNA sequences is often within the range tested here (between 45 and 85 bp; [2]). As expected, estimates of contamination improve with more informative sites, particularly for low contamination rates.

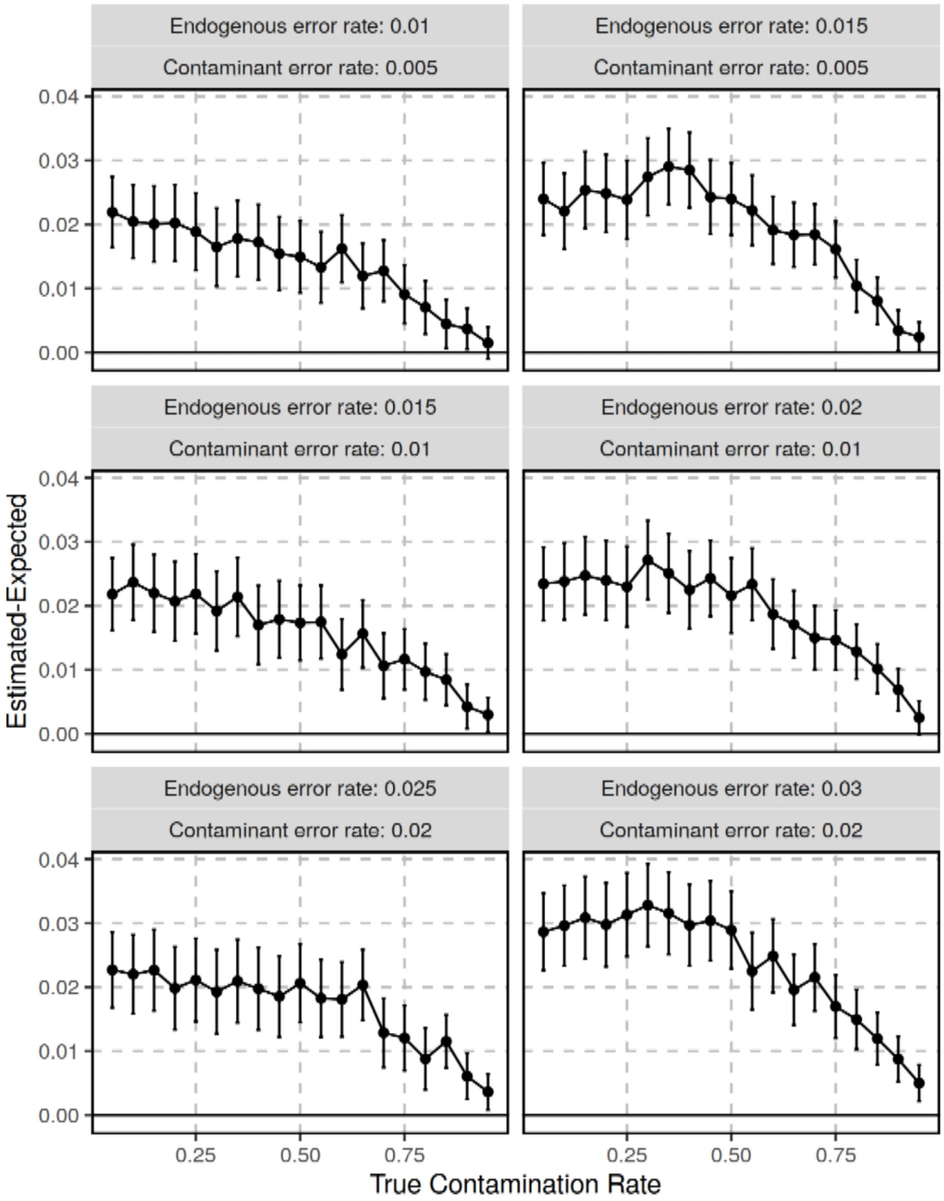

**Figure S2 : Effect of different error (including polymorphism) rates between the endogenous and contaminating sequences.** We simulated sequences with and without deamination-induced substitutions as described for the tests presented in **Figure 4**, but used different error (and polymorphism) rates to challenge our assumption that these should not differ. We used a higher rate of substitutions for the endogenous sequences because the ancient genome will be more distantly related to the reference genome than the present-day contaminant in most cases. The rates are reported on top of each facet. We then mixed these sequences in different proportions for a total of 100,000 sequences and estimated the proportions with AuthentiCT. The y-axis corresponds to the difference between the estimated and expected contamination proportions. Bars represent 95% confidence intervals. In all tested conditions, the very high rate of substitutions on the ancient sequences lead to overestimates of contamination.

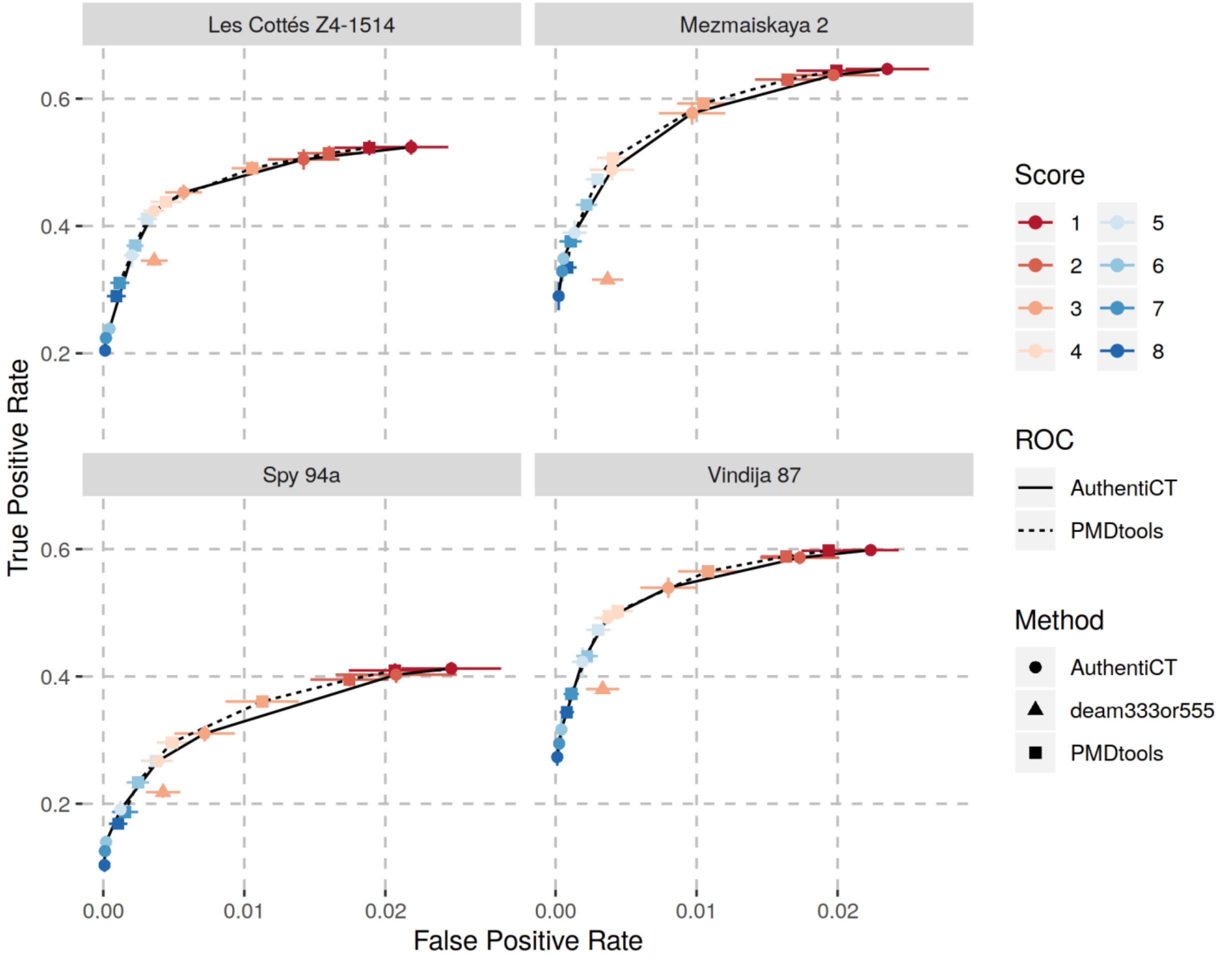

**Figure S3 : Comparison of AuthentiCT and PMDtools to classify ancient and present-day DNA sequences for multiple datasets (when false positive rates are below 0.03).** Each facet corresponds to the comparison for a different Neandertal dataset (random sequences from published datasets, i.e. pools of different libraries, mixed with different proportions of present-day human DNA sequences). For more details, see the caption of **Figure 6**. Note that some additional present-day human DNA sequences may be among the Neandertal sequences: 0.18% (95% Confidence Interval (CI): 0.00-0.42%) for Les Cottes Z4-1514, 0.83% (95% CI: 0.00-1.52%) for Mezmaiskaya 2, 1.75% (95% CI: 0.58-2.84%) for Spy 94a and 1.15% (95% CI: 0-2.37%) for Vindija 87 [3]. However, these low contamination proportions should have a negligible effect on the true positive rates reported given the observed low false positive rate of both methods. Finally, based on the results for these datasets, we would recommend using a score of 4 which maximizes the true positive rate while the false positive rate remains similar to the one observed for the filter of sequences with signs of ancient DNA damage (at least one C-to-T substitution within the first and last three positions, “deam333or555”).

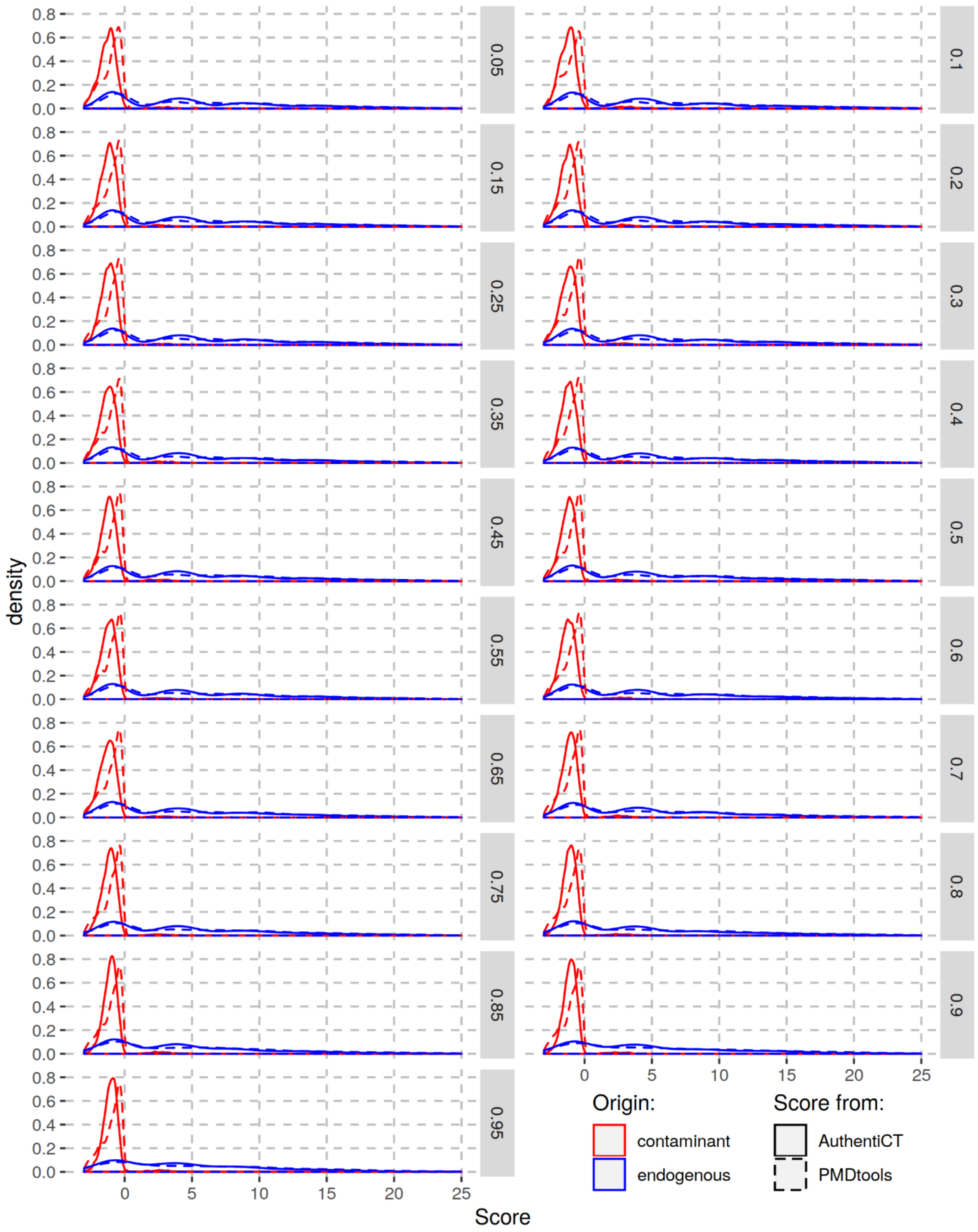

**Figure S4 : Comparison of the distributions of likelihood ratios («Score») computed with AuthentiCT (solid lines) and PMDtools (dashed lines) for different mixtures of ancient and present-day DNA sequences.** Each facet represent a mixture with a different proportion of present-day DNA sequences (human control) and ancient sequences (Mezmaiskaya 2). The scores for the contaminant and ancient sequences are ploted in red and blue, respectively. Note the difference for the red distributions.

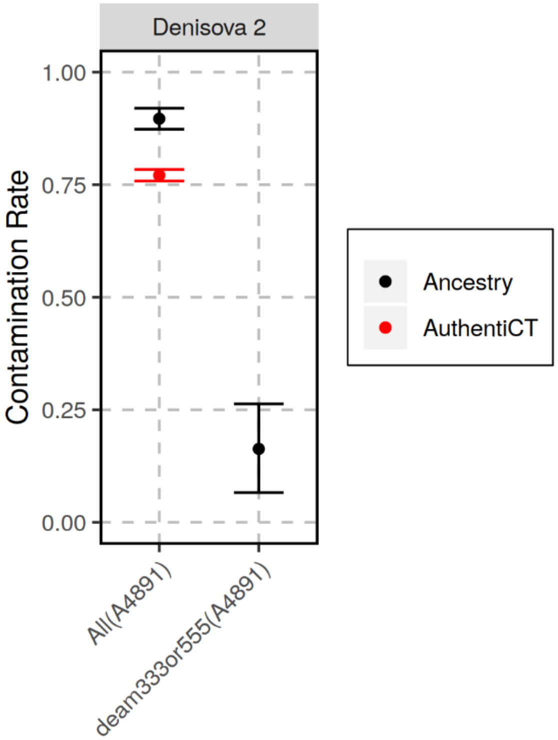

**Figure S5 : Example of a Denisovan sample (Denisova 2, [4]) with a likely deaminated contaminant and the associated contamination underestimate from AuthentiCT.** We used sequences from library A4891 (sequences longer than 35bp and with a mapping quality of at least 25) and estimated contamination from all sequences ("All") or from sequences with one or more C-to-T substitutions within the first or last three positions ("deam333or555"). We used both AuthentiCT and the method based on the allele sharing with a present-day human genome (Results, [5]), although we cannot apply AuthentiCT to the subset of putatively deaminated sequences only. Even after filtering for sequences with C-to-T substitutions, we estimate that 16.3% [95% Confidence Interval: 6.6-26.3%] of sequences still represent modern human DNA, suggesting that the contaminant is deaminated.

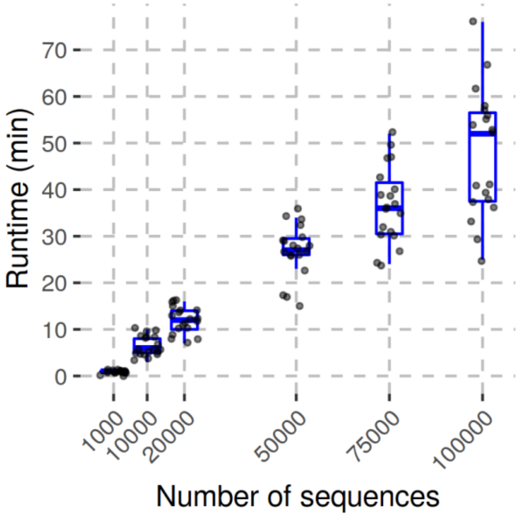

**Figure S6 : Runtime (wall clock) to estimate contamination depending on the number of sequences.** We used 19 artificial mixtures of sequences from a Neandertal dataset (Vindija 33.19) and the present-day human control in varying proportions. We down sampled these datasets to the different number of sequences reported on the x-axis. We used a tolerance of $10^{-10}$ to decide when the maximization of the likelihood should stop. All 114 runs were done in parallel on 114 cores of a multi-core Intel Xeon server (144 CPUs, 1200 MHz each, with a total of 512GB of RAM).

### Tables

**Table S1 : Contamination estimates in the Neandertal datasets used in Figure 3.** The ancestry-based estimates were computed from the proportion of shared alleles with a present-day human (Results, [5]). AuthentiCT was applied to 10,000 sequences that overlap the informative positions used for the ancestry-based estimates. The estimates reported in the second column correspond to the published estimates for the complete datasets [3, 6]. We only used sequences of at least 35bp and with a mapping quality of at least 25.

| Specimen | Contamination Estimate for the complete dataset | Library ID used in **Figure 3** | Ancestry-based | AuthentiCT |
|---|---|---|---|---|
| Les Cottés Z4-1514 | 0.18% [0.00-0.42%] | A9290 | 0 [0.0-0.031] | 0.001 [0-0.031] |
| Mezmaiskaya 2 | 0.83% [0.00-1.52%] | R1917 | 0.011 [0.0-0.088] | 0.015 [0-0.033] |
| Spy 94a | 1.75% [0.58-2.84%] | A9416 | 0 [0.0-0.061] | 0.006 [0-0.056] |
| Vindija 33.19 | 0.21% [0.18-0.23%] | A9369 | 0 [0.0-0.035] | 0.001 [0-0.040] |

**Table S2 : Summary of the libraries used to compare the contamination estimates from AuthentiCT and a method based on the proportion of shared derived alleles with a present-day human genome («Ancestry-based» estimates).** In the third column, we report whether the estimates from AuthentiCT in **Figure 5** were based on sequences overlapping the sites used for the ancestry-based contamination estimates (« info ») or any sequence (« all », if there were not enough sequences to estimate contamination). The fourth column correspond to the number of sequences used for these estimates. Both estimates for all sequences or the ones overlapping informative sites are reported in the last two columns. The numbers in brackets correspond to the 95% confidence intervals. For most libraries, we used sequences of at least 35bp. However, we used a threshold of 30bp for the libraries from the Hohlenstein-Stadel (HST) and Scladina Neandertals to maximize the number of sequences [5]. We also used different length thresholds for the data generated from the Sima hominins (46, 32, 34, 35 and 35bp for the femurXIII, femur fragment, incisor, molar and scapula, respectively, [7, 8]) to reduce the rate of spurious alignments of microbial DNA [7]. In all cases, we used sequences with a mapping quality of at least 25. Mez1: Mezmaiskaya 1; Mez2: Mezmaiskaya 2; NA: Not Applicable.

| Cave / Specimen | Library ID | Data | Number of seq. | Ancestry-based | AuthentiCT (info) | AuthentiCT (all) |
|---|---|---|---|---|---|---|
| Goyet | A9229 | info | 10,000 | 0.03 [0.0-0.106] | 0.097 [0.060-0.134] | 0.070 [0.034-0.106] |
| | A9349 | info | 10,000 | 0 [0.0-0.063] | 0.090 [0.053-0.127] | 0.067 [0.032-0.103] |
| HST | F7016 | info | 10,000 | 0.222 [0.143-0.303] | 0.250 [0.225-0.274] | 0.177 [0.153-0.202] |
| | F7017 | info | 10,000 | 0.184 [0.105-0.265] | 0.254 [0.229-0.279] | 0.184 [0.159-0.210] |
| | L5264 | info | 1,956 | 0.217 [0.041-0.399] | 0.160 [0.110-0.210] | 0.094 [0.070-0.118] |
| | L5265 | info | 1,083 | 0.515 [0.271-0.773] | 0.395 [0.301-0.488] | 0.289 [0.258-0.320] |
| | L5386 | info | 10,000 | 0.173 [0.095-0.253] | 0.220 [0.195-0.244] | 0.163 [0.138-0.188] |
| | R5784 | info | 1,078 | 0.182 [0.0-0.425] | 0.309 [0.234-0.384] | 0.216 [0.191-0.241] |
| | R5785 | info | 1,730 | 0.437 [0.244-0.637] | 0.260 [0.190-0.331] | 0.167 [0.140-0.194] |
| Les Cottes | A9230 | info | 10,000 | 0.011 [0.0-0.087] | 0.001 [0-0.032] | 0.001 [0-0.029] |
| | A9290 | info | 10,000 | 0 [0.0-0.031] | 0.001 [0-0.031] | 0.001 [0-0.030] |
| Mez1 | R5661 | info | 10,000 | 0 [0.0-0.031] | 0.033 [0.003-0.063] | 0.042 [0.014-0.070] |
| | R5662 | info | 10,000 | 0.057 [0.0-0.134] | 0.059 [0.031-0.088] | 0.064 [0.039-0.090] |
| Mez2 | R1917 | info | 10,000 | 0.011 [0.0-0.088] | 0.015 [0-0.033] | 0.001 [0-0.020] |
| | R1916 | all | 8,609 | 0 [0.0-0.5] | NA | 0.014 [0-0.035] |
| Scladina | F7490 | info | 4,575 | 0.654 [0.53-0.781] | 0.627 [0.597-0.657] | 0.498 [0.473-0.523] |
| | G2159 | info | 4,743 | 0.618 [0.498-0.741] | 0.646 [0.616-0.675] | 0.519 [0.496-0.542] |
| | G2160 | info | 5,543 | 0.678 [0.565-0.793] | 0.643 [0.615-0.671] | 0.518 [0.494-0.541] |
| | G2161 | info | 5,404 | 0.683 [0.569-0.8] | 0.633 [0.603-0.664] | 0.528 [0.505-0.551] |
| Sima | femurXIII | all | 10,000 | 0.99 [0.82-0.99] | NA | 0.835 [0.799-0.872] |
| | femur frag. | info | 2,613 | 0.773 [0.607-0.946] | 0.660 [0.615-0.705] | 0.573 [0.545-0.602] |
| | incisor | info | 6,896 | 0.784 [0.682-0.889] | 0.840 [0.822-0.859] | 0.763 [0.745-0.781] |
| | molar | all | 10,000 | 0.82 [0.49-0.99] | 0.850 [0.793-0.907] | 0.741 [0.712-0.769] |
| | scapula | all | 10,000 | 0.881 [0.646-0.999] | NA | 0.927 [0.906-0.949] |
| Spy | A9336 | all | 8,954 | 0.38 [0.0-0.99] | NA | 0.473 [0.439-0.507] |
| | A9416 | info | 10,000 | 0 [0.0-0.061] | 0.006 [0-0.056] | 0.001 [0-0.048] |
| | R5556 | info | 10,000 | 0.052 [0.0-0.129] | 0.001 [0-0.051] | 0.026 [0-0.069] |
| Vindija33.19 | A9368 | info | 10,000 | 0 [0.0-0.035] | 0.014 [0-0.052] | 0.021 [0-0.059] |
| | A9369 | info | 10,000 | 0 [0.0-0.035] | 0.001 [0-0.040] | 0.001 [0-0.041] |
| Vindija87 | A9228 | info | 10,000 | 0 [0.0-0.069] | 0.001 [0-0.020] | 0.005 [0-0.023] |
| | A9348 | info | 10,000 | 0 [0.0-0.057] | 0.001 [0-0.020] | 0.001 [0-0.019] |

#### Supplementary Note 1: Excess of adjacent C-to-T substitutions inside ancient DNA sequences

We found an excess of adjacent C-to-T substitutions in the internal part of sequences compared to expectations from a geometric distribution. Here, we give more details about this comparison.

We first identified sequences with two C-to-T substitutions, masking the first and last five positions as they might fall in single-stranded overhangs. From the first C-to-T substitution (the closest one to the 5' end) in these sequences, we then recorded the distance to the next C-to-T substitution (toward the 3' end). We compared this distribution of distances to a geometric distribution with parameter corresponding to the empirically observed rate of C-to-T substitutions downstream of the first C-to-T substitution (masking also the last five positions). This rate should represent the probability that the next position is a C-to-T substitution if positions are independent. We computed these distributions for sequences from archaic humans and a present-day human control [7]. While we see an excess of adjacent C-to-T substitutions for most datasets, this is not the case for the sequences of the present-day human control. To compare observed and expected distributions, we performed a chi-square goodness-of-fit test. This was done only for the first ten frequencies of these distributions (distances from 1 to 10) as expected counts get too low for higher distances. The p-values are lower than $10^{-15}$ for the archaic datasets, but equal to 0.1967 for the control. We note that the results are similar if we mask the first and last ten positions to identify sequences with two C-to-T substitutions and to compute the rate of C-to-Ts (**Figure S7**).

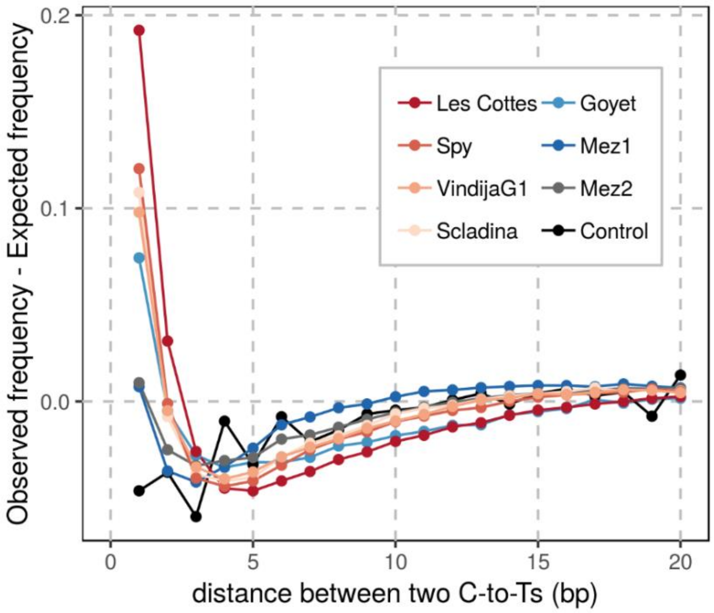

**Figure S7 : Comparison of the observed and expected distributions of distances between C-to-T substitutions.** In contrast to **Figure 1**, we masked the first and last ten positions. See the caption of **Figure 1** for more details.

#### Supplementary Note 2: Inference of the structure of ancient DNA fragments

Damage-associated substitutions along ancient DNA sequences provide some information to reconstruct the structure of the corresponding ancient DNA fragments. In this section, we evaluate with simulations the ability of our Hidden Markov Model (HMM) to infer the location of single-stranded and double-stranded regions.

We simulated ancient DNA sequences using different length distributions and GC contents, but kept the error rates (0.002) and rates of cytosine deamination in single-stranded (0.87) and double-stranded regions (0.015) constant (the other parameters were set to $o$=0.5, $o_2$=0.5, $l_o$=0.34, $l_{ss}$=0.20 and $l_{ds}$=0.003). Using our HMM, we then computed the posterior probability for each state at every informative base along the sequences. The length distributions and the GC contents did not have much influence on the sensitivity and specificity of the classification of regions. In this optimal setting, where the underlying structure of ancient DNA fragments is generated from the same model used for inference, the identifiability of the different regions is nearly perfect, except for the internal single-stranded regions. For this state, the true positive rate does not exceed 0.73 for false positive rates under 0.02 (**Figure S8**). Although a single C-to-T substitution at a terminal position is highly informative about the presence of a single-stranded overhang, a C-to-T substitution toward the middle of an ancient DNA sequence provides limited information to determine whether it happened in a single-stranded or double-stranded context.

We then explored how the rates of substitutions (from deamination or sequencing errors and polymorphisms) influence the power to infer the DNA structure. We only tested error/polymorphism rates up to 10% as most highly divergent sequences will be discarded during alignment [9]. The rate of sequencing errors and polymorphisms also had little influence on the performance to distinguish single-stranded from double-stranded regions (**Figure S9**). In contrast, a lower rate of deamination-induced substitutions decreased the sensitivity (lower positive rate) and specificity (higher false positive rate) of the classification, particularly for internal single-stranded regions (**Figure S10**).

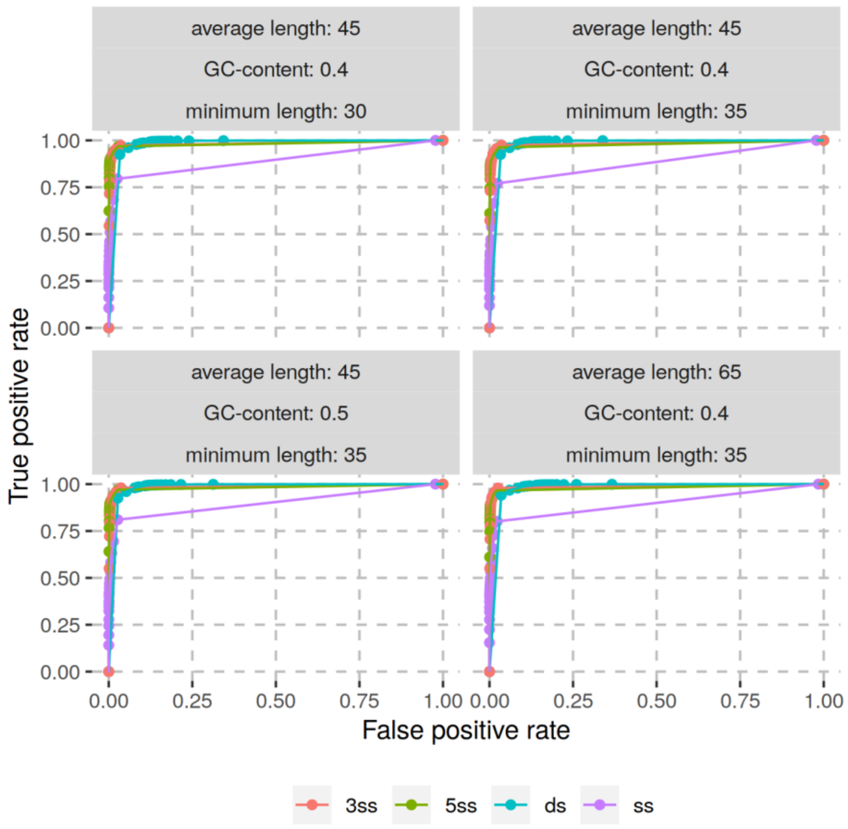

**Figure S8 : Performance of the HMM to infer the structure of simulated ancient DNA fragments based on C-to-T substitutions along the corresponding sequences.** We show the Receiver Operating Characteristic (ROC; with the true positive rate on the y-axis and the false positive rate on the x-axis) curves for the four different hidden states (different colours). Dots represent the results for different cut-offs on the posterior probability used for classification. The facets correspond to different simulated datasets where we varied the GC content and the minimum and average length of sequences.

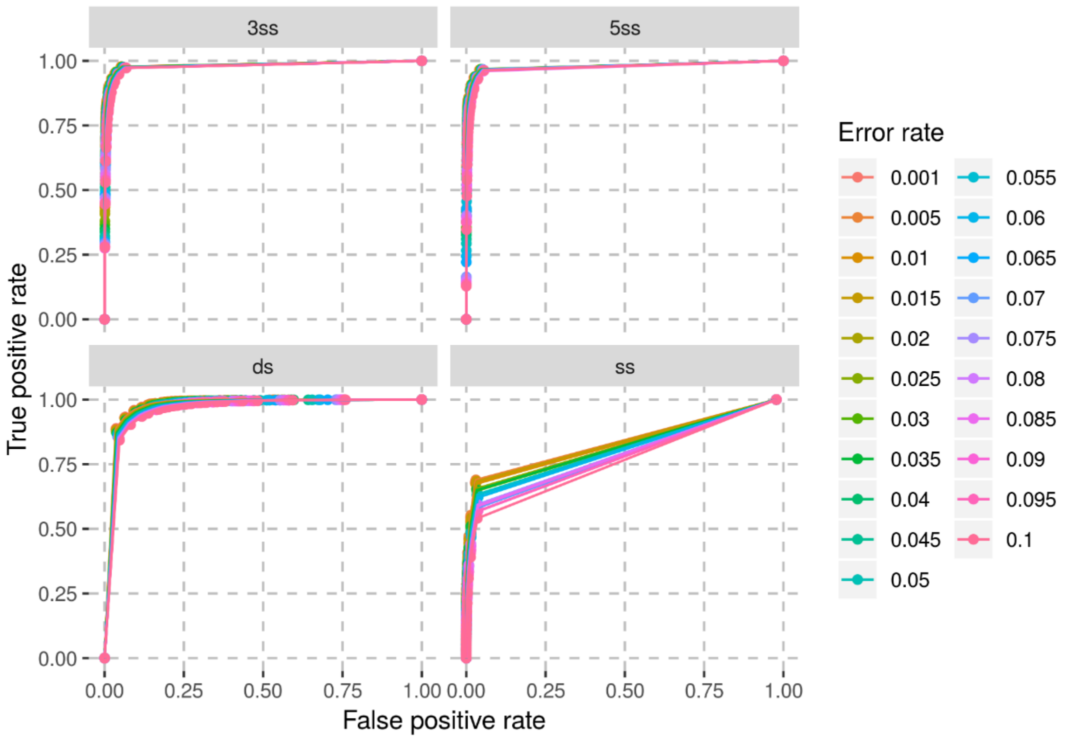

**Figure S9 : Performance of the HMM to infer the structure of simulated ancient DNA fragments with different substitution rates to the reference genome.** The results for the four hidden states are shown in the different facets. The different colours of the ROC curves represent the different substitution rates used in the simulations. The error rate corresponds to the rate of substitutions due to both sequencing errors and polymorphisms.

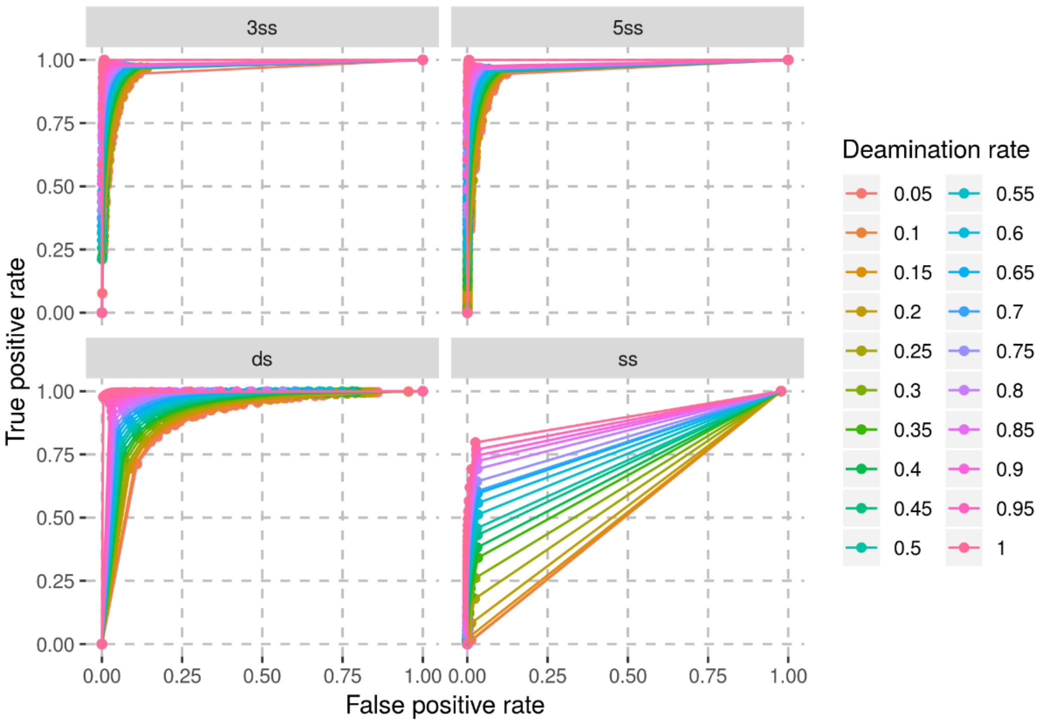

**Figure S10 : Performance of the HMM to infer the structure of simulated ancient DNA fragments with different deamination rates.** The different facets show the true and false positive rates for the four hidden states. The different colours of the ROC curves correspond to the different deamination rates used in the simulations. The deamination rate is the rate of C-to-T substitutions in the single-stranded state.

#### Supplementary Note 3: Differences in contamination estimates per sequence and per base

Contamination estimates from AuthentiCT are, by design, estimates of the proportion of sequences that represent present-day DNA contamination. However, for most downstream analyses, estimates per base are more useful. This is relevant as contamination estimates per sequence and per base may differ if the length of contaminant and endogenous DNA fragments follow different distributions. This is the case for some datasets used in this study. For example, in both datasets from the Hohlenstein-Stadel and Scladina Neandertals, sequences with at least one C-to-T substitution within the first or last three positions are shorter than average (**Figure S11**). Other Neandertal datasets do not show this pattern (**Figure S12**), which suggests the presence of a population of longer contaminating sequences in the datasets of the Hohlenstein-Stadel and Scladina Neandertals. In addition, contamination estimates increase with sequence length, further supporting this hypothesis (**Figure S13**).

These length differences could explain why contamination estimates from AuthentiCT are lower than estimated from the proportion of shared derived alleles with a present-day human genome (see Material and Methods for details about this method) for several libraries prepared from these specimens (**Table S2**). In the main text, we ran AuthentiCT with sequences overlapping the set of positions used for the other method to get contamination estimates that we can compare (except for four libraries with not enough sequences overlapping the informative sites; see **Table S2** for estimates from both analyses, and the number and type of sequences used for each library). This approach lead to more similar estimates between the methods.

In conclusion, we recommend estimating contamination on multiple subsets of the data to check whether contamination estimates per sequence and per base differ. We note that it is preferable to use sequences overlapping the same sites that are later used for the analyses, but one could use sequences overlapping any positions chosen at random to get estimates per base.

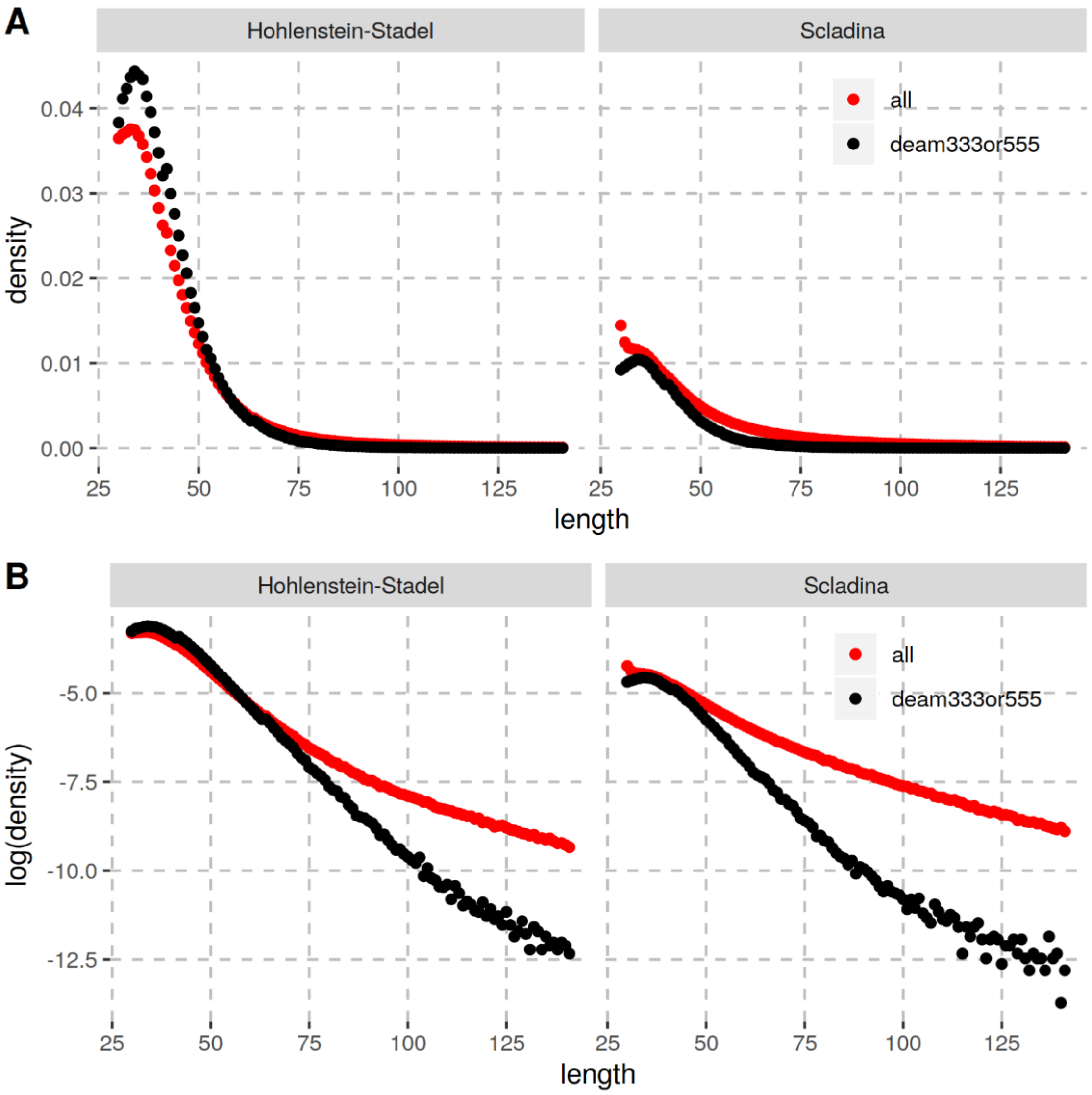

**Figure S11 : Differences in the sequence length distributions among sequences of the Hohlenstein-Stadel and Scladina Neandertals depending on the presence or not of putatively deaminated bases.** We compared the length distribution between all sequences (red dots) and only those that carry at least one C-to-T substitution to the human reference genome within the first or last three positions (black dots). Each facet represents a different Neandertal dataset. Panel **(A)** shows the raw distributions, whereas panel **(B)** shows the logarithm transformation of these distributions to magnify the differences. If a length distribution followed a simple exponential distribution, the logarithm transformation should lead to a linear relationship between the number and length of sequences. Instead, the observed curves may suggest a more complex distribution, potentially a mixture of sequences from present-day humans and from the Neandertal. More importantly, the higher number of long sequences among all sequences compared to the deaminated ones suggests that the present-day human DNA fragments are on average longer than the endogenous Neandertal DNA fragments.

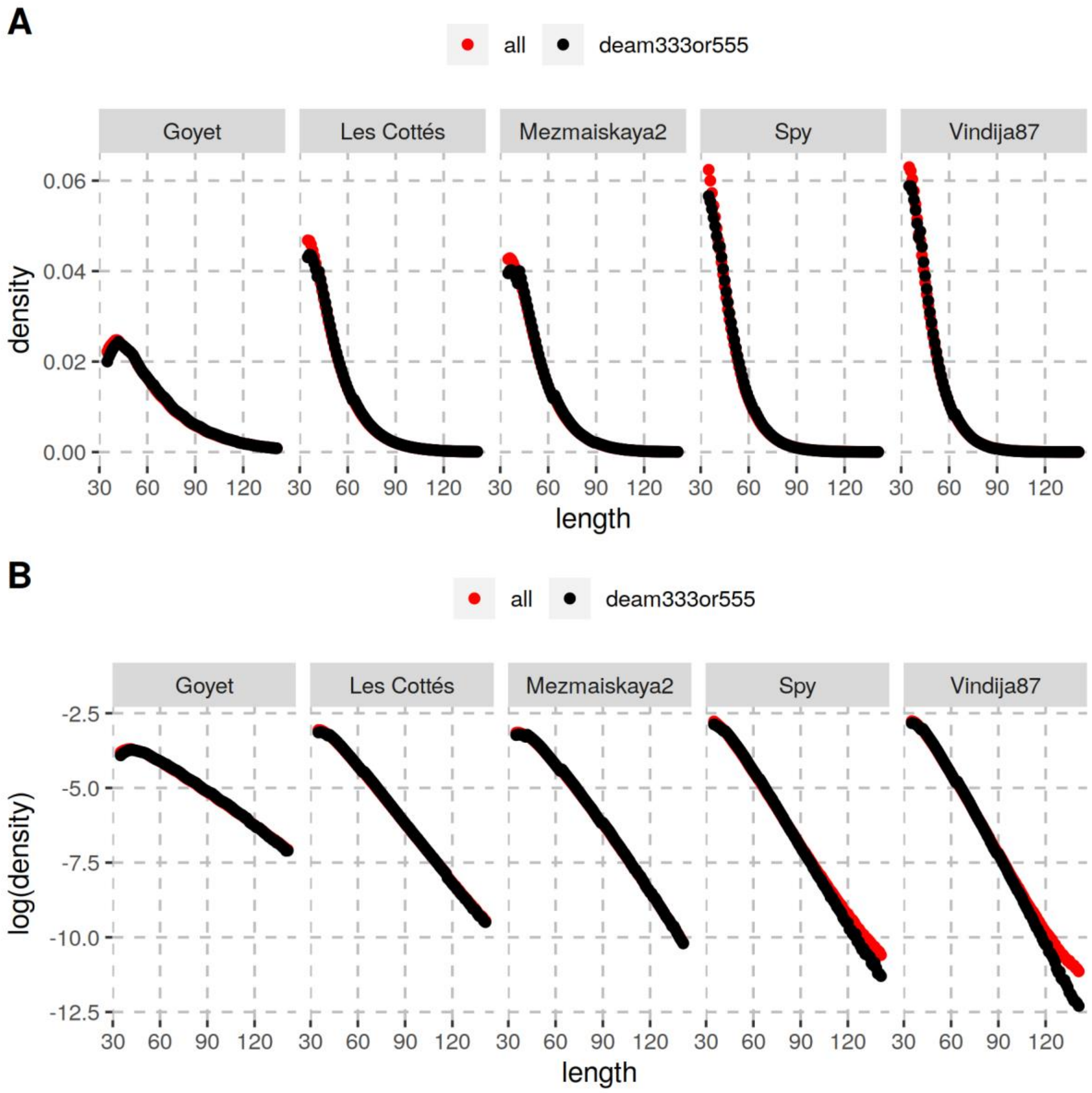

**Figure S12 : Differences in the sequence length distributions among sequences of several Neandertals depending on the presence or not of putatively deaminated bases.** See **Figure S11** for more details and for comparison.

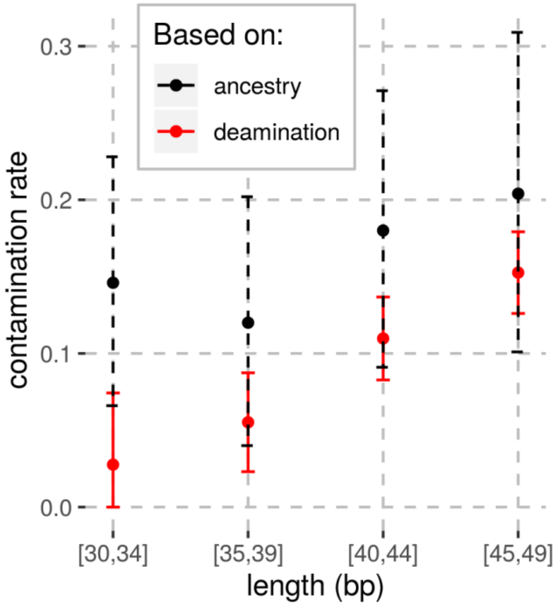

**Figure S13 : Estimates of present-day DNA contamination depending on sequence length in the Hohlenstein-Stadel Neandertal sequencing data.** Red dots represent estimates from AuthentiCT, while black dots represent the estimates obtained from the proportion of shared derived alleles with an African genome at positions where this African genome is different from a Neandertal, a Denisovan and 4 Great Apes (as done previously, [5]). The bars correspond to the 95% confidence intervals.
